# Exploring the effects of CO_2_ elevation on seedlings’ growth of Acacia senegal in the era of changes

**DOI:** 10.1101/2021.09.14.460402

**Authors:** Abdelmoniem A. Attaelmanan, Ahmed A. H. Siddig, Essam I. Warrag

**Affiliations:** University of Khartoum, Faculty of Forestry, Khartoum North postal code 13314, Sudan; Ministry of Social Development; Dept. Environmental Conservation – University of Massachusetts Amherst, USA; Africa Relation Center, Northwest Agriculture & Forestry University, Africa relation Centre, China

**Keywords:** *Acacia senegal*, climate change, drylands, elevated CO_2_, Savanna, Sudan

## Abstract

*Acacia senegal* is a priority and important C_3_ tree species in drylands of Sudan and across the gum belt. Investigation of its seedlings response to elevated carbon dioxide (eCO_2_) is important as atmospheric ([CO_2_]) has increased and predicted to continue to rise. Many studies showed that eCO_2_ causes increased photosynthesis in plants, which leads to greater production of carbohydrates and biomass, and increased soil organic matter and carbon content. This study investigated the effects of eCO_2_ on *A. senegal* seedlings grown in sand and silt soils under irrigation intervals of every day and every two days. Seven days old seedlings were assigned to the treatments in Split - spilt plot design for 4 weeks. The main plot is eCO_2_ (600-800 ppm) and ambient (≤400 ppm) under Free Air CO_2_ Enrichment (FACE) system. Subplots are irrigation intervals and soil types. Seedling height and number of leaves were measured weekly, and seedlings were harvested after 4 weeks where growth parameters and soil properties were measured. The eCO_2_ showed no effect on the measured parameters except the significant increase in tap-root length. However, the irrigation every day showed significant increase than every two days in seedling’s height, number of leaves, root length and seedling’s dry weight but not seedling’s and soil C% & N%. Soil treatment showed effects on stem height, leaf number, seedling’s dry weight, leaves and root N% and soil C% but not root length, seedling C% and soil N%. The results indicate the importance of soil moisture, physical and chemical properties that reflects adaptation of the species to its dry land environment.

## 1. INTRODUCTION

Atmospheric carbon dioxide (CO_2_) emission and concentration has risen since the start of the industrial revolution [1] and continued to rise [2–4]. Trees and plants may respond to rising CO_2_ as elevated atmospheric (eCO_2_) acts to increase photosynthetic activity [5], plant growth and plant biomass production of many plant species [6,7]. Drylands vegetation include C_3_ and C_4_ trees, shrubs and grasses [8]. In some of eCO_2_ experiments, C_4_ plants showed little or no enhancement of growth (dry matter production) in contrast, C_3_ species showed 3 times of that experienced by C_4_ plants in stimulation of photosynthesis by eCO_2_ [9].

The acacias are important C_3_ dryland species [10]. *A. senegal* is a multi-purpose tree producing gum Arabic a high-value export commodity from Sudan and some African countries, and important component of traditional dryland agroforestry resilience system and source of livelihoods in the Sudan [11]. The tree also provides animal fodder, multiple timber products, intercropping, firewood, food and medicines [12,13]. Furthermore, it is one of the most important sub-Saharan African trees inhabiting Savanna systems that are under threat of ongoing anthropogenic and climate-mediated degradation and that have led to substantial losses of natural habitats [14].

Seedlings are most responsive to eCO_2_ where the early growth enhancement under eCO_2_ accelerates ontogeny and pattern of growth [15,16]. The effect of eCO_2_ on acacias as important components of the dryland natural plant communities’ needs to be studied. Soil types and soil moisture content are important determinants to *A. senegal* response [17]. The methodological and experimental developments such as the Free-Air Carbon Enrichment (FACE) are effective way to quantify the effects of eCO_2_ on trees in field settings [18,19]. The FACE designs provide means for studies without any direct perturbation of microclimate [20]. Therefore, the objective of the paper is to experimentally examine the responses of *A. senegal* seedlings to eCO_2_ under varying water and soil conditions using the FACE design.

## 2. MATERIALS AND METHODS

### 2.1. Study site & settings

The experiment was conducted in the nursery as Split-split plot design with the CO_2_ treatments as main plots, and the water interval and soil types as subplots. The nursery of the Faculty of Forestry, University of Khartoum at Shambat, Khartoum North-Sudan (15° 40’ 5” North, 32 32’ 1” East). Shambat has a subtropical desert / low-latitude arid hot climate. The experiment was conducted using FACE system for eCO_2_ in the range of 600 to 800 and ambient treatments. The watering intervals were every day and every two days, while the soil treatments are sand and silt. Bulk seeds collected from El-Damazeen forests was obtained from the National Tree Seed Center and germinated in polymer bags of 10×20cm filled with silt or sand soils) and irrigated daily. After one week 60 seedlings per soil type were selected with minimum morphological variation (i.e. almost same size & branching pattern) among them. They were then assigned randomly in the experimental plots. The plot was divided into 4 subplots of five seedlings. Sixty seedlings of equal size from each soil type were assigned randomly to irrigation treatments. Three pairs of 1m×2m plots were prepared and each one assigned randomly for the eCO_2_ and the other for the ambient CO_2_ treatments. Then each plot was divided into two parts of 1×1m, and assigned randomly for the irrigation treatments every day and every two days. Then 5 seedlings from those raised in silt soil and 5 ones from sandy soil were assigned randomly to each of the watering treatments (total of 120 seedlings, 60 silt and 60 sand).

### 2.2. Measured variables

Seedlings’ height was measured and new leaves were counted weekly. After four weeks the plants were harvested, length of the tap root was measured and dry weight of the leaves, stems and roots was weighed for each plan separately. Seedling’s C% and N% were determined using CHNS-O Analyzer and applying Standard Test Method (ASTM International, model D 5291-02. 2002, USA) for instrumental determination of carbon and nitrogen of plant.

Soil C% measurement is based on the oxidation of organic C with dichromate in acid medium [21] and soil N% was measured by using Bremner’s method [22].

### 2.3. Data analysis

The Analysis of Variance (ANOVA) procedures and Duncan’s Multiple Range Test to separate means of the same factor at significance were carried out using SAS. The model is Split-split plot with three blocks, CO_2_ (main plot), watering interval and soil type within the watering interval. The model used in the experiment was:

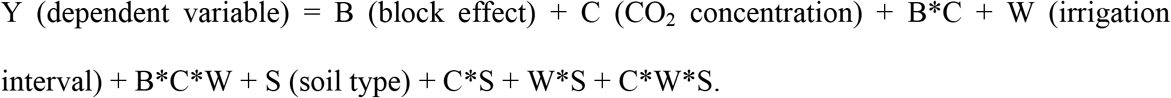

## 3. RESULTS

### 3.1. Effects of eCO_2_ concentration on seedlings’ growth parameters

The eCO_2_ concentration had no effect on seedling’s height, number of leaves per seedling after 1 week, 2 weeks, 3 weeks and 4 weeks from the start of the experiment. In addition to leaves, stem, root, total dry weight, C% and N% were not affected (Table 1). Similarly, eCO_2_ had not significantly affected soil media C%, N% and C/N. While tap root length was significantly affected by CO_2_ elevation (P=0.002) as eCO_2_ had decreased tap root length by 16% (P≤ 0.05; Table 1).

**Table 1:**
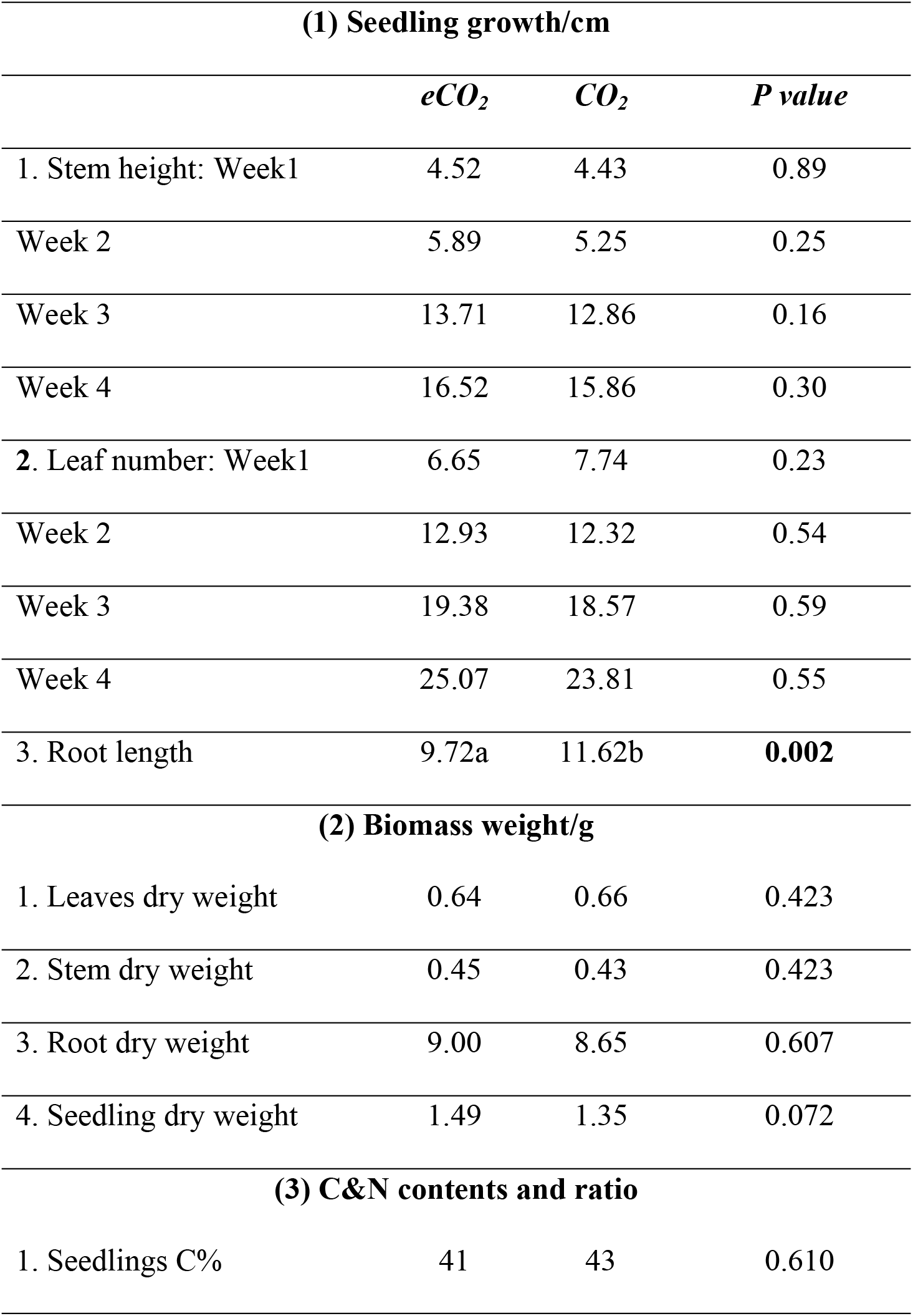
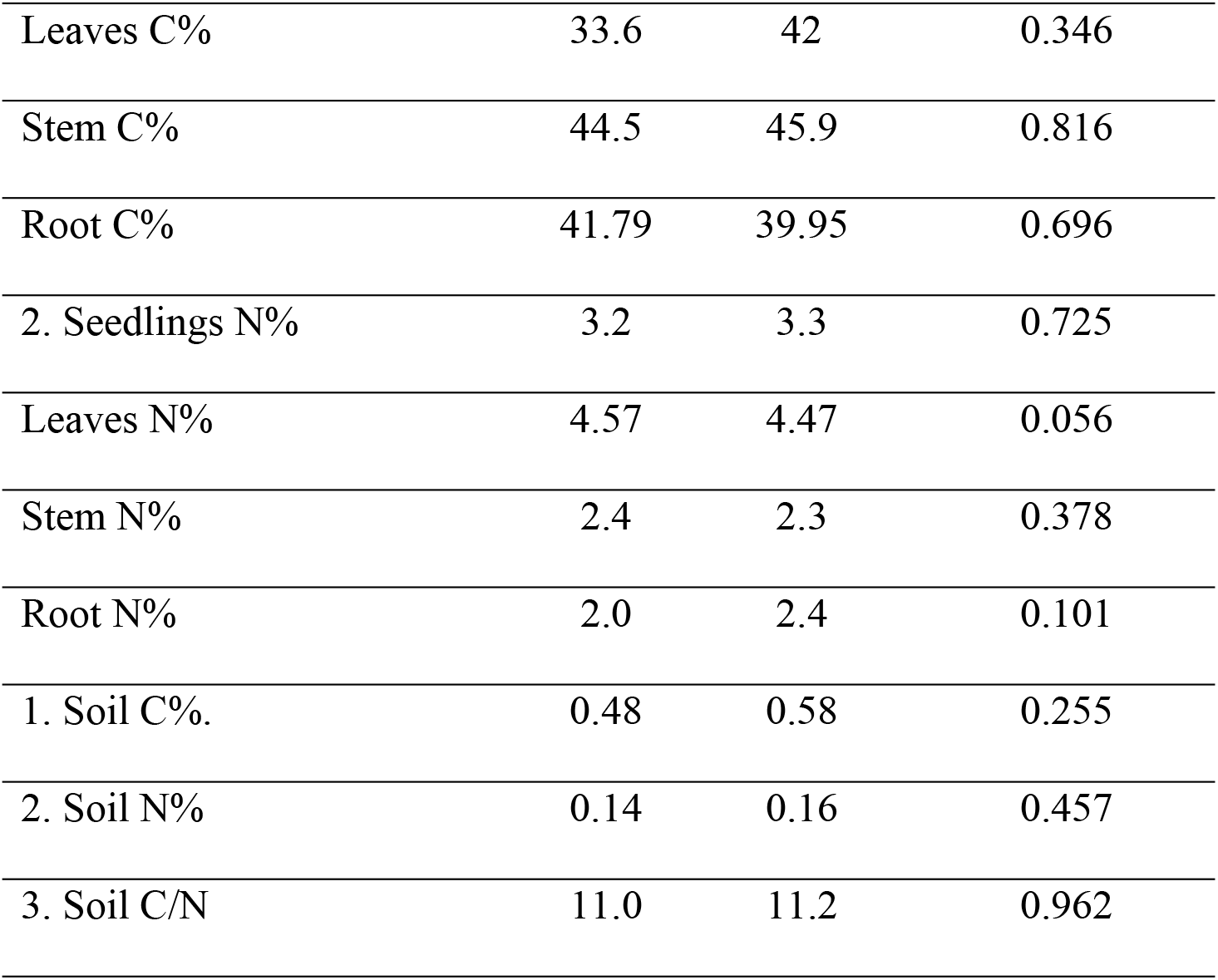
Effects of CO_2_ concentration on seedling’s height, number of leaves, root length, seedling’s biomass weight and seedling and soil carbon & nitrogen contents after 4 weeks.

| **(1) Seedling growth/cm** |  |  |  |
| --- | --- | --- | --- |
|  | *eCO<sub>2</sub>* | *CO<sub>2</sub>* | *P value* |
| 1. Stem height: Week1 | 4.52 | 4.43 | 0.89 |
| Week 2 | 5.89 | 5.25 | 0.25 |
| Week 3 | 13.71 | 12.86 | 0.16 |
| Week 4 | 16.52 | 15.86 | 0.30 |
| 2. Leaf number: Week1 | 6.65 | 7.74 | 0.23 |
| Week 2 | 12.93 | 12.32 | 0.54 |
| Week 3 | 19.38 | 18.57 | 0.59 |
| Week 4 | 25.07 | 23.81 | 0.55 |
| 3. Root length | 9.72a | 11.62b | **0.002** |
| **(2) Biomass weight/g** |  |  |  |
| 1. Leaves dry weight | 0.64 | 0.66 | 0.423 |
| 2. Stem dry weight | 0.45 | 0.43 | 0.423 |
| 3. Root dry weight | 9.00 | 8.65 | 0.607 |
| 4. Seedling dry weight | 1.49 | 1.35 | 0.072 |
| **(3) C&N contents and ratio** |  |  |  |
| 1. Seedlings C% | 41 | 43 | 0.610 |

|  |  |  |  |  |
| --- | --- | --- | --- | --- |
| 326 | Leaves C% | 33.6 | 42 | 0.346 |
| 327 |  |  |  |  |
| 328 | Stem C% | 44.5 | 45.9 | 0.816 |
| 329 |  |  |  |  |
| 330 | Root C% | 41.79 | 39.95 | 0.696 |
| 331 |  |  |  |  |
| 332 | 2. Seedlings N% | 3.2 | 3.3 | 0.725 |
| 333 |  |  |  |  |
| 334 | Leaves N% | 4.57 | 4.47 | 0.056 |
| 335 |  |  |  |  |
| 336 | Stem N% | 2.4 | 2.3 | 0.378 |
| 337 |  |  |  |  |
| 338 | Root N% | 2.0 | 2.4 | 0.101 |
| 339 |  |  |  |  |
| 340 | 1. Soil C%. | 0.48 | 0.58 | 0.255 |
| 341 |  |  |  |  |
| 342 | 2. Soil N% | 0.14 | 0.16 | 0.457 |
| 343 |  |  |  |  |
| 344 | 3. Soil C/N | 11.0 | 11.2 | 0.962 |
| 345 |  |  |  |  |
| 346 |  |  |  |  |
| 347 |  |  |  |  |
| 348 |  |  |  |  |
| 349 |  |  |  |  |
| 350 |  |  |  |  |
| 351 |  |  |  |  |
| 352 |  |  |  |  |
| 353 |  |  |  |  |
| 354 |  |  |  |  |
| 355 |  |  |  |  |
| 356 |  |  |  |  |
| 357 |  |  |  |  |
| 358 |  |  |  |  |
| 359 |  |  |  |  |
| 360 |  |  |  |  |
| 361 |  |  |  |  |
| 362 |  |  |  |  |
| 363 |  |  |  |  |
| 364 |  |  |  |  |
| 365 |  |  |  |  |
| 366 |  |  |  |  |
| 367 |  |  |  |  |
| 368 |  |  |  |  |
| 369 |  |  |  |  |

### 3.2. Effects of irrigation interval on seedlings’ growth parameters

Irrigation interval had significant effect on seedling’s height, leaf number and tap root length at the end of the first, second, third and fourth week (p≤ 0.050 -< 0.0001; Table 2). Irrigation every day had resulted in increase in seedling’s height by 42%, 36%, 37% and 46% by the end of first, second, third and fourth weeks, respectively. Also, It increased number of leaves per seedling at the end of each week (45, 40, 37 and 61% respectively). Similarly irrigation every day had increased tap root length by 24% at the end of fourth week (p≤ 0.05; Table 2).

**Table 2:**
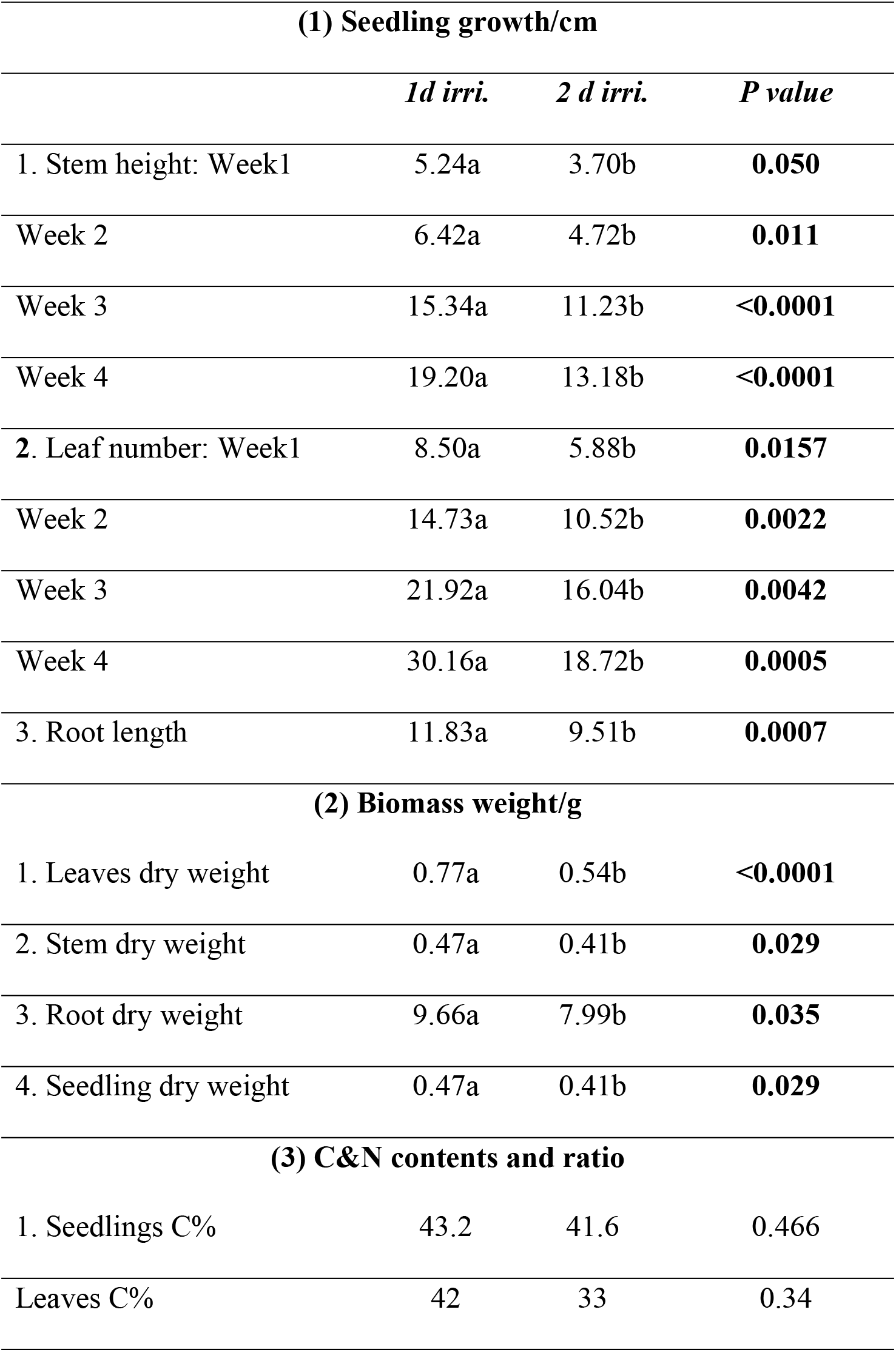
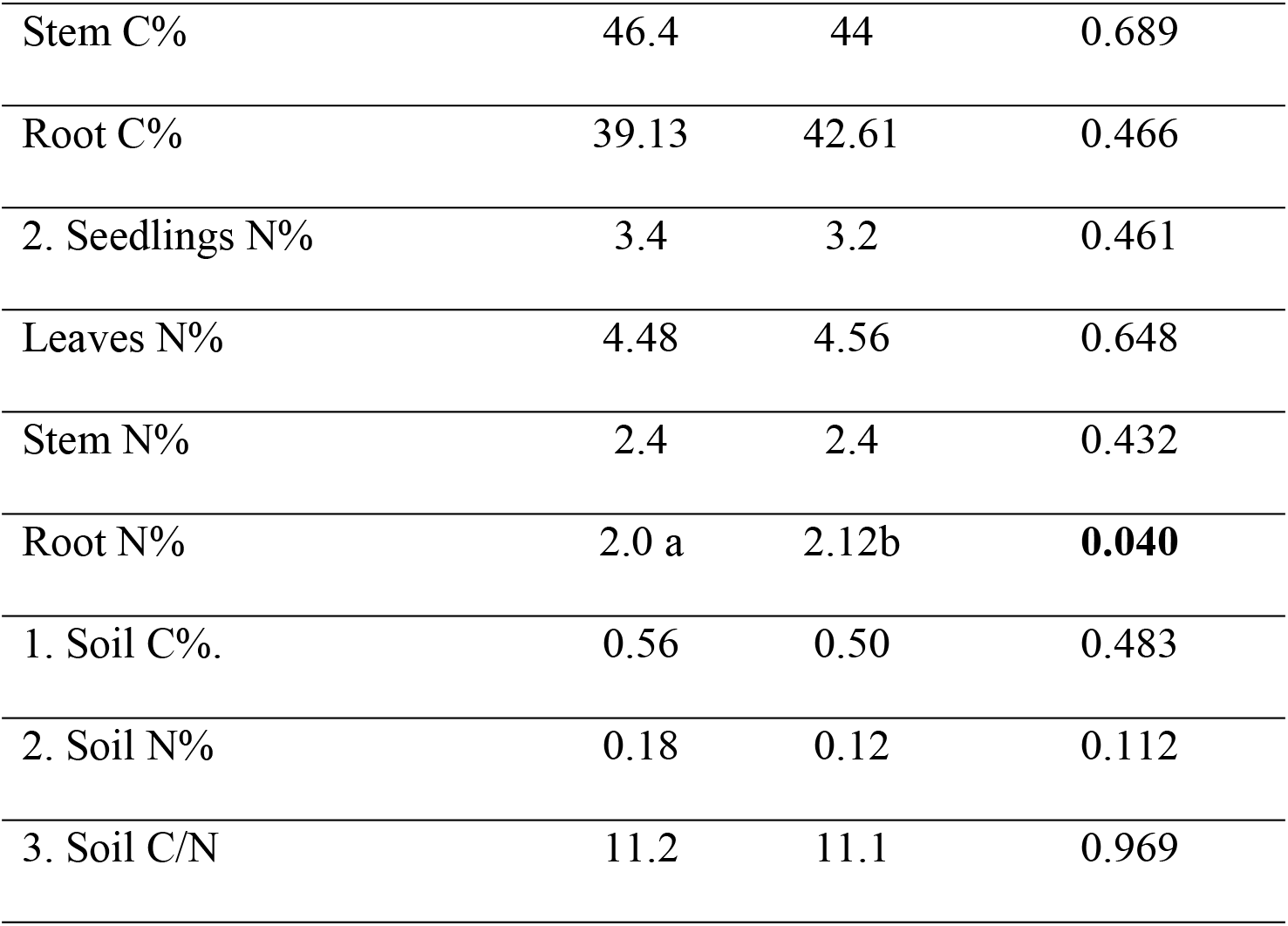
Effects of irrigation interval on seedling’s height, number of leaves, root length, seedling’s biomass weight and seedling & soil carbon and nitrogen contents after 4 weeks.

| **(1) Seedling growth/cm** |  |  |  |
| --- | --- | --- | --- |
|  | *1d irri.* | *2 d irri.* | *P value* |
| 1. Stem height: Week1 | 5.24a | 3.70b | **0.050** |
| Week 2 | 6.42a | 4.72b | **0.011** |
| Week 3 | 15.34a | 11.23b | **<0.0001** |
| Week 4 | 19.20a | 13.18b | **<0.0001** |
| 2. Leaf number: Week1 | 8.50a | 5.88b | **0.0157** |
| Week 2 | 14.73a | 10.52b | **0.0022** |
| Week 3 | 21.92a | 16.04b | **0.0042** |
| Week 4 | 30.16a | 18.72b | **0.0005** |
| 3. Root length | 11.83a | 9.51b | **0.0007** |
| **(2) Biomass weight/g** |  |  |  |
| 1. Leaves dry weight | 0.77a | 0.54b | **<0.0001** |
| 2. Stem dry weight | 0.47a | 0.41b | **0.029** |
| 3. Root dry weight | 9.66a | 7.99b | **0.035** |
| 4. Seedling dry weight | 0.47a | 0.41b | **0.029** |
| **(3) C&N contents and ratio** |  |  |  |
| 1. Seedlings C% | 43.2 | 41.6 | 0.466 |
| Leaves C% | 42 | 33 | 0.34 |

|  |  |  |  |  |
| --- | --- | --- | --- | --- |
| 373 | Stem C% | 46.4 | 44 | 0.689 |
| 374 |  |  |  |  |
| 375 | Root C% | 39.13 | 42.61 | 0.466 |
| 376 |  |  |  |  |
| 377 | 2. Seedlings N% | 3.4 | 3.2 | 0.461 |
| 378 |  |  |  |  |
| 379 | Leaves N% | 4.48 | 4.56 | 0.648 |
| 380 |  |  |  |  |
| 381 | Stem N% | 2.4 | 2.4 | 0.432 |
| 382 |  |  |  |  |
| 383 | Root N% | 2.0 a | 2.12b | **0.040** |
| 384 |  |  |  |  |
| 385 | 1. Soil C%. | 0.56 | 0.50 | 0.483 |
| 386 |  |  |  |  |
| 387 | 2. Soil N% | 0.18 | 0.12 | 0.112 |
| 388 |  |  |  |  |
| 389 | 3. Soil C/N | 11.2 | 11.1 | 0.969 |
| 390 |  |  |  |  |
| 391 |  |  |  |  |
| 392 |  |  |  |  |
| 393 |  |  |  |  |
| 394 |  |  |  |  |
| 395 |  |  |  |  |
| 396 |  |  |  |  |
| 397 |  |  |  |  |
| 398 |  |  |  |  |
| 399 |  |  |  |  |
| 400 |  |  |  |  |
| 401 |  |  |  |  |
| 402 |  |  |  |  |
| 403 |  |  |  |  |
| 404 |  |  |  |  |
| 405 |  |  |  |  |
| 406 |  |  |  |  |
| 407 |  |  |  |  |
| 408 |  |  |  |  |
| 409 |  |  |  |  |
| 410 |  |  |  |  |
| 411 |  |  |  |  |
| 412 |  |  |  |  |
| 413 |  |  |  |  |
| 414 |  |  |  |  |
| 415 |  |  |  |  |
| 416 |  |  |  |  |
| 417 |  |  |  |  |

Irrigation interval had affected leaves, stem, root and seedling weights (P ≤ 0.035-0.0001). Irrigation every day, as compared to irrigation every two days, had increased weight of leaves by 43%, stem by 15%, root by 21% and seedling by15% (p≤ 0.05; Table 2).

Irrigation interval had not affected seedling’s C%, N% and its compartments, soil C%, N% and soil C/N. Except root N% was affected (P=0.040) as irrigation every day had decreased root N by 6%. However, the daily irrigation was numerically lower C/N than two days irrigation but not significant (p≤ 0.05; Table 2).

### 3.3. Effects of soil type on seedlings’ growth parameters

Soil type showed significant effects on seedling’s height (first, third and fourth week) and leaf number (second, third and fourth week) but not length of tap root (P=0.027-<0.0001; Table 3). The silt soil had increased seedling’s height by 66% for the first week, by 29% for the third and by 31% for the fourth week. Also, silt soil had enhanced leaf number by 22% for the second, by 26% for the third and by 33% for the fourth week (P≤ 0.05; Table 3). Silt soil gave longer root (9%) but not significant.

**Table 3:**
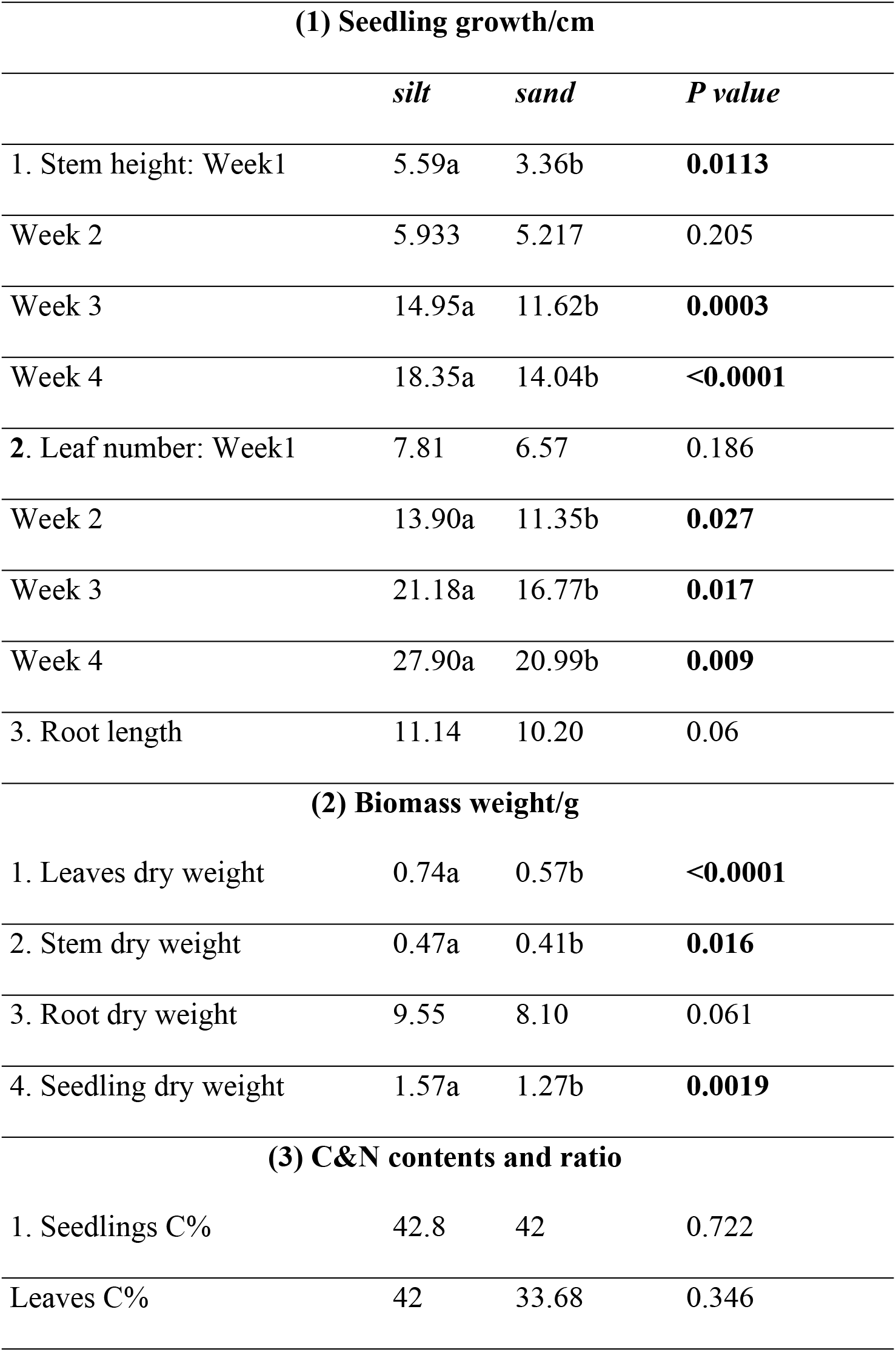
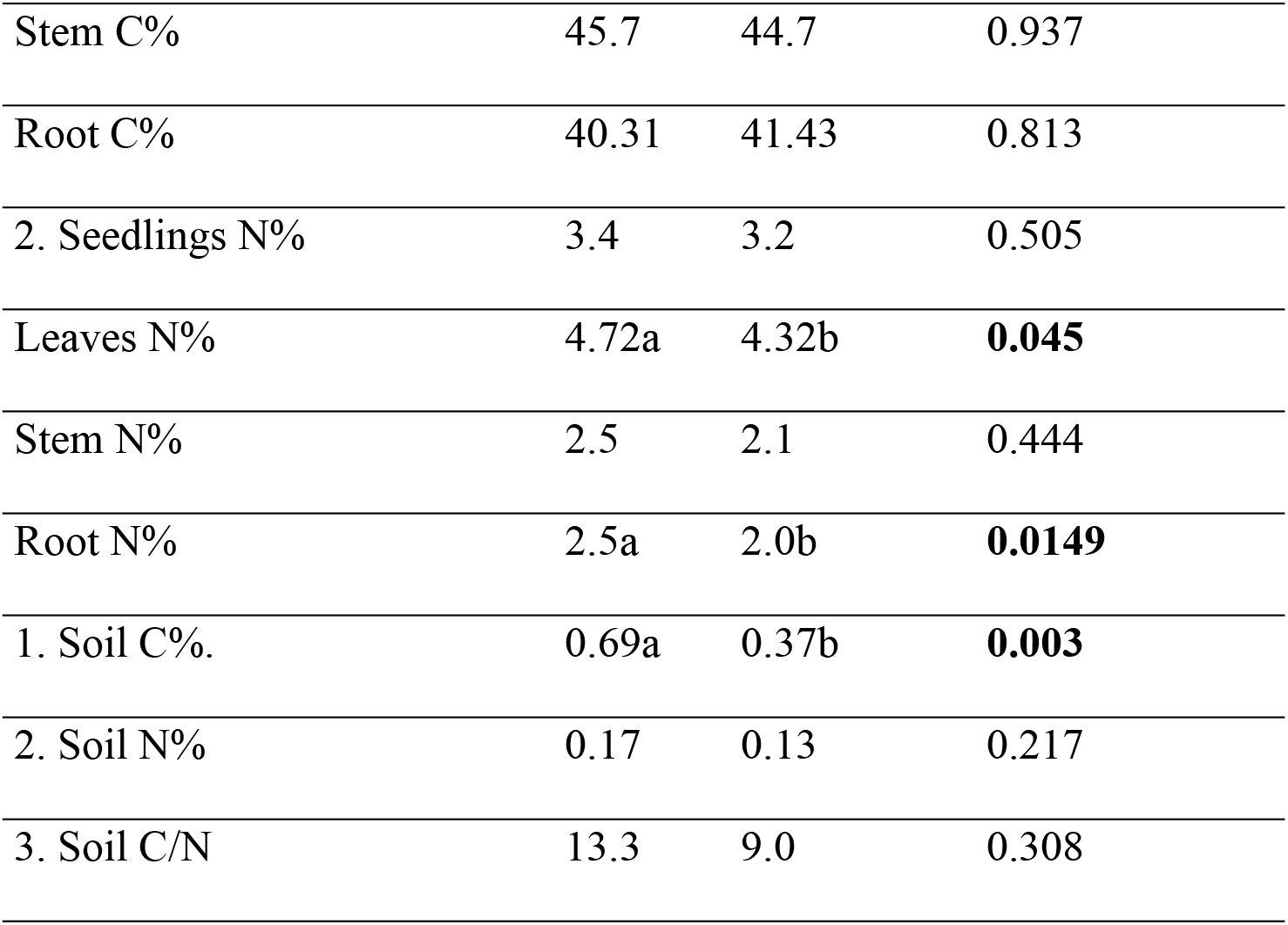
Effects of soil type on seedling’s height, number of leaves, root length, seedling’s biomass weight and seedling & soil carbon & nitrogen contents after 4 weeks.

| **(1) Seedling growth/cm** |  |  |  |
| --- | --- | --- | --- |
|  | *silt* | *sand* | *P value* |
| 1. Stem height: Week1 | 5.59a | 3.36b | **0.0113** |
| Week 2 | 5.933 | 5.217 | 0.205 |
| Week 3 | 14.95a | 11.62b | **0.0003** |
| Week 4 | 18.35a | 14.04b | **<0.0001** |
| 2. Leaf number: Week1 | 7.81 | 6.57 | 0.186 |
| Week 2 | 13.90a | 11.35b | **0.027** |
| Week 3 | 21.18a | 16.77b | **0.017** |
| Week 4 | 27.90a | 20.99b | **0.009** |
| 3. Root length | 11.14 | 10.20 | 0.06 |
| **(2) Biomass weight/g** |  |  |  |
| 1. Leaves dry weight | 0.74a | 0.57b | **<0.0001** |
| 2. Stem dry weight | 0.47a | 0.41b | **0.016** |
| 3. Root dry weight | 9.55 | 8.10 | 0.061 |
| 4. Seedling dry weight | 1.57a | 1.27b | **0.0019** |
| **(3) C&N contents and ratio** |  |  |  |
| 1. Seedlings C% | 42.8 | 42 | 0.722 |
| Leaves C% | 42 | 33.68 | 0.346 |

|  |  |  |  |  |
| --- | --- | --- | --- | --- |
| 421 | Stem C% | 45.7 | 44.7 | 0.937 |
| 422 |  |  |  |  |
| 423 | Root C% | 40.31 | 41.43 | 0.813 |
| 424 |  |  |  |  |
| 425 | 2. Seedlings N% | 3.4 | 3.2 | 0.505 |
| 426 |  |  |  |  |
| 427 | Leaves N% | 4.72a | 4.32b | **0.045** |
| 428 |  |  |  |  |
| 429 | Stem N% | 2.5 | 2.1 | 0.444 |
| 430 |  |  |  |  |
| 431 | Root N% | 2.5a | 2.0b | **0.0149** |
| 432 |  |  |  |  |
| 433 | 1. Soil C%. | 0.69a | 0.37b | **0.003** |
| 434 |  |  |  |  |
| 435 | 2. Soil N% | 0.17 | 0.13 | 0.217 |
| 436 |  |  |  |  |
| 437 | 3. Soil C/N | 13.3 | 9.0 | 0.308 |
| 438 |  |  |  |  |
| 439 |  |  |  |  |
| 440 |  |  |  |  |
| 441 |  |  |  |  |
| 442 |  |  |  |  |
| 443 |  |  |  |  |
| 444 |  |  |  |  |

Soil type had significant effects of leaves’ dry weight (P <0.0001), stem (P=0.016) and seedling (P=0.0019) but it had slightly insignificant effects on root’s dry weight (P=0.061). Consequently, silt soil had higher dry weights in leaves by 30%, stem by 15% and seeding by 24% (P≤ 0.05; Table 3).

Soil type had no effects on seedling’s, leaves’, stem’s and root’s C and N content. Similarly, soil N% and C/N were not affected by soil type, but soil C% was affected by soil type (P=0.003). However, the N% and C/N of silt were numerically higher than sand soil.

## 4. DISCUSSION

### 4.1. Effects of eCO_2_ concentration on growth parameters

The insignificancy of eCO_2_ on growth parameters (height, leaf number, root length, seedling dry weight and seedling C%, N%) is in line with many other literatures [23–25]. On the other hand, eCO_2_ is known to increase plant growth productivity and consequently has stimulated overall forests biomass growth [26–29]. Such discrepancy might be due to differences in genetic characteristics of the studied species, duration of studies, CO_2_ elevation (exposure) techniques, and sites microclimates. Our results can be explained as, less photosynthetic machinery that made by N deficiency or other nutrients causing lower rates of photosynthesis resulted in no increasing in net primary productivity (NPP).

### 4.2. Effects of eCO_2_ concentration on soil carbon and nitrogen content

The irresponsive of soil C%, N% and C/N to eCO_2_ is in agreement with various studies [30,31]. On the other hand, the results contrast the findings of [32,33]. The difference of these results can be attributed to the adverse response through down regulation of photosynthesis when plants exposed to higher CO_2_ concentrations beyond the certain thresholds, or the rapid rate of CO_2_ assimilation requires correspondingly other nutrients specially foliar N which is experienced to be declined under elevation CO_2_.

## 5. CONCLUSION

The irresponsive of most measured variables of *A. senegal* to eCO_2_ concentration and the high significant effects of water and soil factors can be attributed to the long time adaptation of the species in drylands of Sudan to water and to some extent soil type. Generally, the growth parameters of the *A. senegal* seedlings were more responsive to the environmentally limiting factors in its natural habitat, such as soil moisture content and soil chemical and physical properties.

Nevertheless, our study was limited in a number of aspects including sample size, duration & design of the experiment. Further studies on incorporating some or all of these factors will give a better picture about responses of *acacias* to eCO_2_ in dryland settings.

According to the results and general trends of this study, it is recommended that:

1. - Water availability is the most important factor of eCO_2_ for seedling growth and hence water harvesting and management will play a key role in the context of elevation of CO_2_.
2. - In drylands where it characterized by low level of precipitation, planting seedling in silt soil is recommended for greatest growth and productivity.
3. - More long-term experiments on *A. senegal* are recommended to evaluate the effects of eCO_2_ for long term periods as some measured variables seems to be affected by time (e.g. biomass).
4. - Other interacted factors with CO_2_ like thermal stress and nutrient limitation should be investigated for better understanding of *Acacias’* response to CO_2_.
5. - The responses of the most important C_3_ tree species in drylands Sudan like *Acacia nilotica, Acacia seyal*,..etc to CO_2_ elevation need to be evaluated for long and short terms.
6. - FACE systems (are being constructed in the USA, and now widely used in other places in the world) can be used sufficiently to study effects of CO_2_ elevation on other trees and crops under natural conditions in Africa.
7. - With regard to the net photosynthesis and stomatal responses it would be advantageous to monitor them under CO_2_ elevation to understand up-regulation or down-regulation of photosynthesis and to show *A. senegal* stomatal response to eCO_2_ and under what conditions this occurs. The responses of stomatal conductance and canopy leaf area to eCO_2_ are important specially in drylands to determine both the short and long-term risk of exposure to drought.

## ACKNOWLEDGEMENTS

This work was supported by Faculty of Forestry, University of Khartoum. I gratefully acknowledge the generous support provided by Dr. Abdelazzim Yaseen and Prof. Abdalla Mirghani El Tayeb for their assistance and encouragement. Special thanks and appreciation are due to Carbon Dioxide Company for their funding and sincere thanks are extended to Mr. Waleed Elshikh, Dr. Waleed Salih, Tawfeg Eltyb Zain, Mawia and Mohammed Nasr for their help and facilitation.

